# Stat3 co-operates with mutant *Ezh2*^Y641F^ to regulate gene expression and limit the anti-tumor immune response in melanoma

**DOI:** 10.1101/2021.06.01.446580

**Authors:** Sarah M. Zimmerman, Samantha J. Nixon, Pei Yu Chen, Leela Raj, Sofia Smith, Rachel L. Paolini, Phyo Nay Lin, George P. Souroullas

## Abstract

One of the most frequently genetically altered chromatin modifiers in melanoma is the Enhancer of Zeste Homolog 2 (*EZH2*), the catalytic component of the Polycomb Repressive Complex 2 (PRC2), which methylates lysine 27 on histone 3 (H3K27me3), a chromatin mark associated with transcriptional repression. Genetic alterations in *EZH2* in melanoma include amplifications and activating point mutations at tyrosine 641 (Y641). The oncogenic role of EZH2 in melanoma has previously been identified; however, its downstream mechanisms remain underexplored. We found that in genetically engineered mouse models, *Ezh2*^Y641F^ expression causes upregulation of a subset of interferon-regulated genes in melanoma cells, suggesting a potential role of the immune system in the pathogenesis of these mutations. Upregulation of these interferon genes was not a result of changes in H3K27me3, but through a direct and non-canonical interaction between Ezh2 and Signal Transducer and Activator of Transcription 3 (Stat3). Ezh2 directly binds Stat3, and, in the presence of Ezh2^Y641F^, Stat3 protein is hypermethylated. Expression of Stat3 was required to maintain an anti-tumor immune response and its depletion resulted in faster melanoma progression and disease recurrence. Molecularly, Stat3 and Ezh2 bind together at many genomic loci, and can both activate and repress gene expression in association with the rest of the PRC2 complex. These results suggest that Ezh2 can mediate melanomagenesis through evasion of the anti-tumor immune response, and that the immunomodulatory properties of Stat3 are context dependent.

## INTRODUCTION

Epigenetic modifiers are frequently mutated in many types of cancers, including melanoma, and are also associated with poor survival, response to treatment, or relapse (1,2). They are thus a relatively new but very promising class of cancer therapeutic targets. Unlike the genetic code, which is static, the epigenome is reversible, creating numerous opportunities for manipulation using small molecule inhibitors to restore a normal epigenetic state or to take advantage of genetic or synthetic lethal interactions (3). The availability of small molecules targeting epigenetic/chromatin modifying enzymes has created an enormous interest in their therapeutic properties, with currently hundreds of ongoing clinical trials testing their efficacy (4,5). Despite clear successes, progress in some areas has been slow, suggesting that we still do not fully understand the downstream mechanisms of these epigenetic modifiers or the interactions between different epigenetic and chromatin modifications. Understanding these mechanisms is, however, an essential element to developing effective therapies.

Epigenetic modifiers regulate fundamental cellular processes such as DNA replication and DNA repair (6), along with many cell-intrinsic hallmarks of cancer such as the cell cycle, cell growth and proliferation. It is thus not surprising that they also play a significant role in the development of cancer. Epigenetic modifiers also contribute towards cell-extrinsic properties relevant to cancer development or progression, such as regulation of various aspects of the immune system (7–9). One of the most frequently genetically altered chromatin modifiers in melanoma is EZH2, the catalytic component of the Polycomb Repressive Complex 2 (PRC2), which methylates lysine 27 on histone 3 (H3K27me3), a chromatin mark associated with transcriptional repression. Specifically, about 20% of melanoma patients in TCGA datasets exhibit either genomic amplifications that include *EZH2*, overexpression of EZH2 protein, or express the activating and neomorphic point mutation *EZH2*^*Y641*^ (**Supp. Fig. 1a, b**). The frequency of *EZH2*^Y641^ mutations ranges from 5-10% of melanoma patients from numerous studies (10–12). Patients with genetic alterations in PRC2 components and other chromatin modifiers that are highly connected with PRC2 exhibit lower survival rates (**Supp. Fig. 1c**), as do patients with high protein expression of EZH2 itself (**Supp. Fig. 1d)**.

Previously, we generated a mouse model permitting conditional expression of *Ezh2*^Y641F^ and showed that *Ezh2*^Y641F^ expression in murine melanocytes caused a global increase and redistribution in H3K27me3 across the genome, while accelerating the onset of melanoma (13). *Ezh2*^Y641F^ in mouse models strongly co-operated with mutant *Braf*^V600E^ but failed to co-operate with mutant *Nras*^Q61R^, two of the main oncogenic drivers in human melanoma (10). This result was consistent with the fact that in human patients, *EZH2*^Y641F^ mutations tend to co-occur with *BRAF*^V600E^ but not *NRAS*^Q61^ mutations. Additionally, both *EZH2* and *BRAF* are located within 100kb on the same chromosomal arm, and chromosomal amplifications tend to include both genes. Despite our understanding of the consequences of the *Ezh2*^Y641F^ mutations on chromatin and distribution of H3K27me3, we still do not know which downstream molecular mechanisms mediated by *Ezh2*^Y641F^ contribute towards melanoma initiation and progression.

In addition to *Ezh2*^Y641F^ mutations, Ezh2 overexpression is also implicated in melanoma development and metastasis (14), suggesting that in general, increased Ezh2 activity in melanocytes may promote transformation through silencing of distinct tumor suppressors (14) or promoting disease progression by controlling mechanisms of adaptive resistance to tumor immunotherapy (15). Melanoma patients have benefited greatly from immunotherapy; however, melanoma remains the deadliest form of skin cancer once it has metastasized, with a considerable number of patients relapsing after targeted treatment, not responding or becoming resistant to checkpoint inhibitors, or experiencing significant toxicity. Therapeutic options at this point become scarce, highlighting our limited understanding of the context and mechanisms of an effective anti-tumor immune response. The epigenome could be an important variable in determining how various immunomodulatory factors affect the anti-tumor immune response.

To understand the oncogenic mechanisms of Ezh2 in melanoma, we analyzed and compared gene expression profiles of *Ezh2*^WT^ vs *Ezh2*^Y641F^ mutant mouse melanoma tumors and identified a unique interferon-related gene signature. Despite the dramatic re-organization of H3K27me3 across the genome mediated by Ezh2^Y641F^, expression of these genes was not dependent on a change in H3K27me3 at their promoters but was still dependent on Ezh2 methyltransferase activity. We thus hypothesized that expression of these genes is driven via non-canonical interactions of Ezh2 with non-histone proteins. One candidate protein was Stat3 which had previously been shown to be methylated by Ezh2 (16,17).

Stat3 is a member of the STAT family and is activated in response to extracellular signaling proteins, including growth factors and cytokines such as IL-6 (18,19). Stat3 is activated by phosphorylation, which promotes its dimerization and transport into the nucleus, where it functions as a transcription factor (20). Stat3 often acts in association with other transcription factors and co-activators (21–23). In addition to being a transcriptional activator, Stat3 can also repress its target genes (24,25); however, the mechanism of how Stat3 represses gene expression is not clear. Stat3 is implicated in many oncogenic mechanisms, contributing to both cell-intrinsic and - extrinsic hallmarks of cancer such as cell growth, proliferation, differentiation, apoptosis, inflammation, and immune cell regulation, and is therefore an attractive therapeutic target in some types of cancer (26).

In this study, we investigated the oncogenic mechanisms of Ezh2 in melanoma, and show that *Ezh2*^Y641F^ in melanoma cells causes consequential changes to the anti-tumor immune response that are dependent on Stat3 expression. We therefore present significant insight into how hyperactive Ezh2 promotes melanoma progression, a possible mechanism for those oncogenic activities, and a potential explanation of how Stat3 suppresses gene expression via interaction with the PRC2 complex.

## MATERIALS & METHODS

### Animals

Animals were housed in an Association for Assessment and Accreditation of Laboratory Animal Care (AAALAC)-accredited facility and treated in accordance with protocols approved by the Institutional Animal Care and Use Committee (IACUC) for animal research at Washington University School of Medicine in St. Louis.

### *In vivo* tumor models

C57Bl/6 female mice were obtained from Taconic Biosciences. Tumor cells were injected subcutaneously in the flank at 0.5 × 10^6 cells per injection, two injections per mouse. Tumor size was within the size permitted by the Washington University IACUC committee. Tumor volume was measured using calipers to measure length and width; volume was calculated using the formula: length × width^2. Anti-CD8 (InVivoMAb anti-mouse CD8β Cat# BE0223 Lot# 733219O1) was administered to the mice intraperitoneally at 10 mg/kg (2.5 mg/ml, 100 µl injection volume). For flow cytometry analysis of infiltrating immune cells, tumors were harvested at Day 9 or, for mice treated with anti-CD8, 15 days after last injection. Tumors were finely chopped, dispersed by passing several times through a syringe with 18G needle, and filtered through a 0.40 µm filter.

### Flow Cytometric analysis

Single cell suspensions from dissociated tumors were washed with PBS contained 1% FBS and stained with antibody cocktails for detection of tumor-infiltrating lymphocytes. The lymphoid antibody cocktail contained: anti-CD45-PerCP/Cy5.5 (BioLegend 103132), anti-NK1.1-FITC (BioLegend 108706), anti-CD3-PB (BioLegend 100214), anti-CD4-APC (BioLegend 100412), anti-CD8-AF700 (BioLegend 100730), and anti-PD-1(CD279)-PE/Cy7 (BioLegend 135216). The myeloid antibody cocktail contained: anti-CD45-PerCP/Cy5.5 (BioLegend 103132), anti-CD19-FITC (BioLegend 115506), anti-B220-FITC (BioLegend 103206), anti-CD3-FITC (BioLegend 100204), anti-CD11b(Mac1)-PB (BioLegend 101224), anti-CD11c-PE/Cy7 (BioLegend 117318), and anti-Ly-6G(Gr1)-AF700 (BioLegend 127622). Cells were stained with propidium iodide to detect dead cells. Samples were run on an Attune Nxt Flow Cytometer (ThermoFisher Scientific) at the Siteman Flow Cytometry Core Facility. Analysis was done in FlowJo V10 and statistically significant differences were identified using One-way ANOVA and Tukey’s post-hoc test performed in R.

### Analysis of RNA-seq data

Gene Set Enrichment Analysis (GSEA) was performed as described here (27). Enrichment of differentially expressed genes was carried out using a pre-ranked gene list using log_2_ ratio of expression of Ezh2^WT^ vs Ezh2^Y641F^. Independent analysis of the RNA-seq data was performed with the Database for Annotation, Visualization, Integrated Discovery (DAVID) functional annotation tools (28). The functional annotation clustering tool was used to analyze gene ontology (GO) terms associated with the differentially expressed genes in Ezh2^Y641F^ cells vs Ezh2^WT^.

### Cell culture, shRNA, cell growth assay

For these studies we used six syngeneic mouse melanoma cell lines expressing *Ezh2*^WT^ (234, 855, and 480; *Tyr*-*Cre*^ERT2^ *Braf*^V600E^ *Pten*^F/+^ *Ezh2*^+/+^) or *Ezh2*^Y641F^ (234Δ, 855Δ, and 480Δ; *Tyr*-*Cre*^ERT2^ *Braf*^V600E^ *Pten*^F/+^ *Ezh2*^Y641F/+^). Cells were cultured in DMEM high glucose (Sigma D6429) with 10% fetal bovine serum (Corning Cat# MT35010CV) and 1% penicillin-streptomycin (Genesee Scientific Cat# 25-512). Stat3 knockdown cell lines were generated by transducing cells with lentiviral shRNA (TRCN0000071456, TRCN0000071454, TRCN0000071453, Sigma). Lentiviruses were generated using 293T cells via transfection with PEI and appropriate vectors, including a VSV-G envelop vector. Cell lines were transduced at an MOI of 1 and puromycin selection (3 µg/ml) was initiated 48 hours after transfection. Knockdown was confirmed by qPCR and immunoblotting. For the growth assay, cells were plated in triplicate in 96-well plates at 500 cells/well and growth was determined by Alamar Blue staining and detection. Cells were incubated in Alamar blue for 1 hour prior to calorimetric measurement. The average number of cells present at each timepoint was compared between cell lines, and statistically significant differences were detected using Student’s t-test.

### Chromatin Immunoprecipitation (ChIP) sequencing and analysis

Ten million cells were used for each ChIP pull down and were prepared using the protocol from Lee et. al. (29). Briefly, cells were lysed and sheared using a probe Sonicator to achieve a size of 200-300 bp, and DNA was isolated using phenol: chloroform for library preparation. Immunoprecipitations were performed overnight at 4°C using anti-Stat3 (Cell Signaling #9139) and anti-Ezh2 (Cell Signaling #5246) antibodies at 1:100 dilution. Libraries were prepared using TruSeq ChIP kit (Illumina) and multiplexed for sequencing using 50-bp single reads. Libraries were sequenced on a Hi-Seq 2000 at the Genome Access Center of Washington University in St. Louis. Processing of ChIP-seq data was carried out using Encode standards. Binding sites were identified using MACS2. Functional significance of Ezh2 and Stat3 binding sites/peaks was evaluated using the Genomic Regions Enrichment of Annotations Tool (GREAT) (30). Data was visualized using the Integrated Genomics viewer (31). De novo motif analysis was carried out using HOMER (32) to find 8-20 bp motifs in the mm10 genome. The motif analysis was done on differential Ezh2 and Stat3 peaks that were higher in the *Ezh2*^WT^ cells, as well as the peaks that were higher in the *Ezh2*^Y641F^ cells to compare the differences. The Ezh2 and Stat3 differentially bound peaks were annotated with the R package ChipSeeker to look at the differences in feature distributions and visualize the genomic annotation of these peaks between *Ezh2*^WT^ and *Ezh2*^Y641F^ cell lines. The python program deeptools was used to generate heatmaps and line plots associated with Ezh2 and Stat3 peaks (33). The plotProfile tool was used to create a profile plot for scores over the genomic regions around the TSS for all the Ezh2 and Stat3 marks.

For ChIP-qPCR and sequential ChIP (ChIP-reChIP), forty million cells per sample were used and prepared according to Furlan-Magaril et al. (34). Quantitative PCR was carried out as described below.

### Protein Immunoprecipitation (IP)

Cells were lysed using lysis buffer containing protease and phosphatase inhibitors on ice and sonicated briefly to complete lysis. Samples were centrifuged for 10 minutes at 16,000 × g at 4°C, and the supernatant was used for IPs. Lysates were incubated with Stat3 antibody (1:100, Cell Signaling #9139) for 24 hours, then incubated with protein A/G magnetic beads (Bio-Rad SureBeads Cat# 1614023) for 1 hour. Samples were eluted by incubating in 1X Laemmli buffer (Bio-Rad Cat# 161-0747) with beta-mercaptoethanol at 70°C for 10 minutes.

### Immunoblotting

Samples were prepared by adding 4X Laemmli buffer with beta-mercaptoethanol (except for IP eluants) and boiled for 5 minutes. Samples were cooled briefly and loaded onto 4-20% pre-cast gels (Bio-Rad Mini-PROTEAN TGX Gels Cat# 4561095) and run at 100-120V using the Bio-Rad Mini-PROTEAN Tetra System. The separated proteins were transferred onto nitrocellulose membranes using the Bio-Rad Trans-Blot Turbo Transfer System at 1.0 Amp for 30 minutes. Membranes were blocked for 1 hour in 5% milk in TBS-T (TBS + 0.1% Tween-20), then incubated with primary antibodies overnight at 4°C. Primary antibodies were: anti-Ezh2 Cell Signaling #5246 at 1:1000, anti-Suz12 Cell Signaling #3737 at 1:1000, anti-Eed Cell Signaling #85322 at 1:1000, anti-beta actin Abcam ab213262 at 1:2000, anti-Stat3 Cell Signaling #9139 at 1:500, and anti-phospho-Stat3 (Tyr705) Cell Signaling #9145 at 1:2000. The membranes were washed with TBS-T before incubating with the secondary antibodies at room temperature for 1 hour in the dark. Secondary antibodies were: anti-rabbit IgG (H+L) (DyLight 800 4X PEG Conjugate) Cell Signaling #5151 at 1:50,000 and anti-mouse IgG (H+L) (DyLight 680 Conjugate) Cell Signaling #5470 at 1:50,000. Membranes were imaged using a Licor Odyssey Infrared Imager, and Licor Image Studio software was used for densitometry analysis.

### Quantitative PCR

RNA was isolated from melanoma cells using the Aurum Total RNA Mini Kit (Bio-Rad Cat# 7326820) according to the manufacturer’s instructions. The igScript First Strand cDNA Synthesis Kit (Intact Genomics Cat# 4314) was used to synthesize the cDNA with the following modification: 1.0 µl of random hexamers plus 1.0 µl of oligo-dT were used as primers for the reverse transcriptase reaction. The cDNA was used undiluted for qPCR with the iTaq Universal SYBR Green Supermix (Bio-Rad Cat# 1725120). Primers used for qPCR are in the table below.

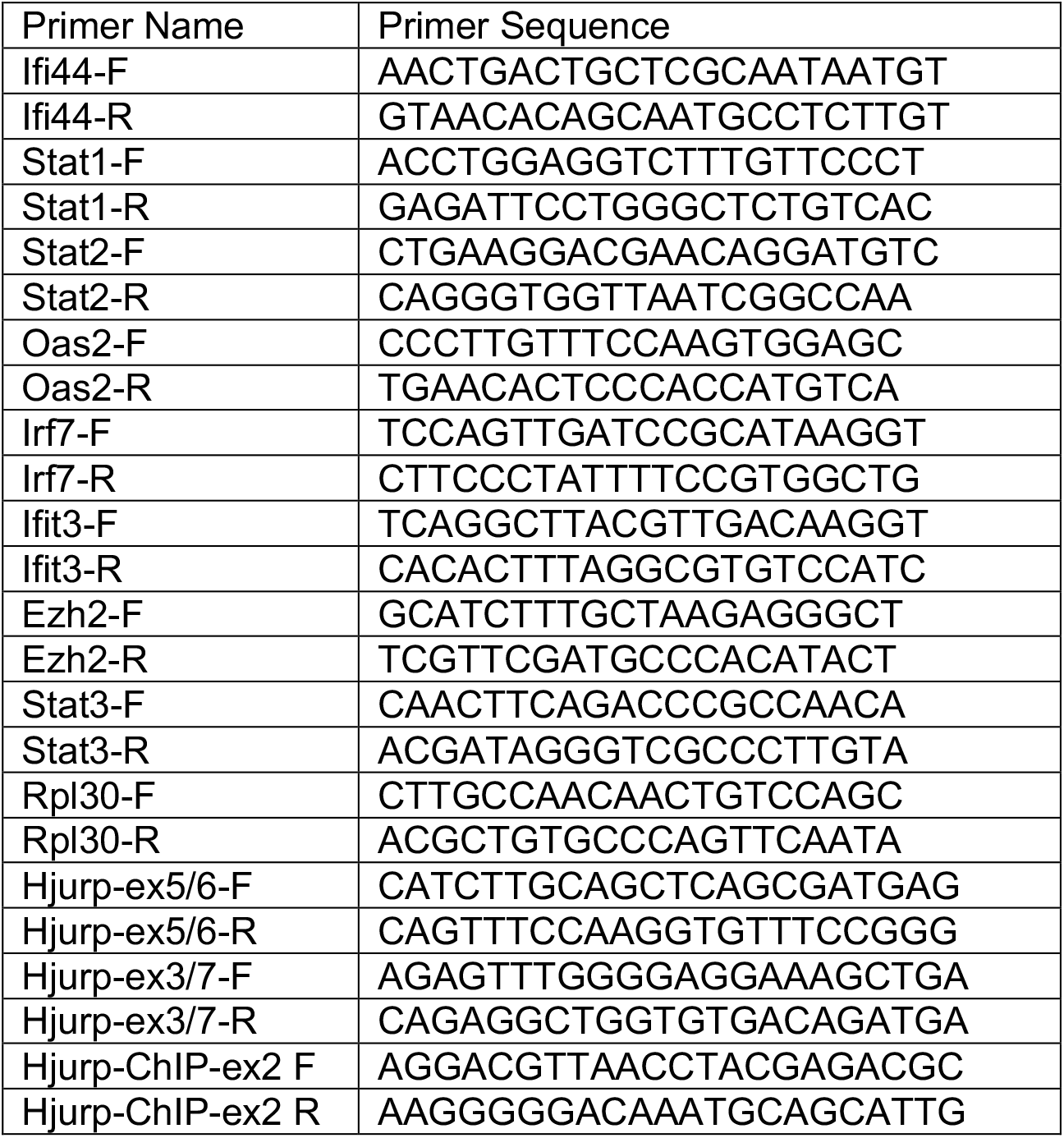

## RESULTS

### Expression of Ezh2^Y641F^ in melanoma induces an interferon-related gene expression signature

In previous studies we generated a mouse allele modeling the *Ezh2*^Y641F^ mutation in melanoma. We found that this allele was a potent driver of melanoma, especially in combination with the *Braf*^V600E^ mutation. Expression of Ezh2^Y641F^ globally increased abundance of H3K27me3, however, it also caused a widespread redistribution of this mark across the genome (13), the mechanisms of which remain unclear. To better understand the underlying oncogenic mechanisms of these mutations, we used isogenic cell lines derived from tumors of *Tyr-Cre*^ERT2^ *Braf*^V600E/+^ *Pten*^F/+^ mice that differ only in expression of the *Ezh2* mutation (WT vs Y641F). Functional annotation (28,35) of differentially expressed genes identified enrichment for several Gene Ontology terms, including proteolysis, cell-cell adhesion, protein post-translational modifications, transcriptional regulation, and immune response (**Fig. 1a**). Gene set enrichment analysis additionally identified Interferon signaling, IFNB1 targets and cytokine signaling as also enriched in *Ezh2*^Y641F^ vs *Ezh2*^WT^ tumors (**Fig. 1b, c**). A subset of these differentially expressed, IFN-related genes was validated on separate biological samples by qPCR (**Fig. 1d**).

**Figure 1.**
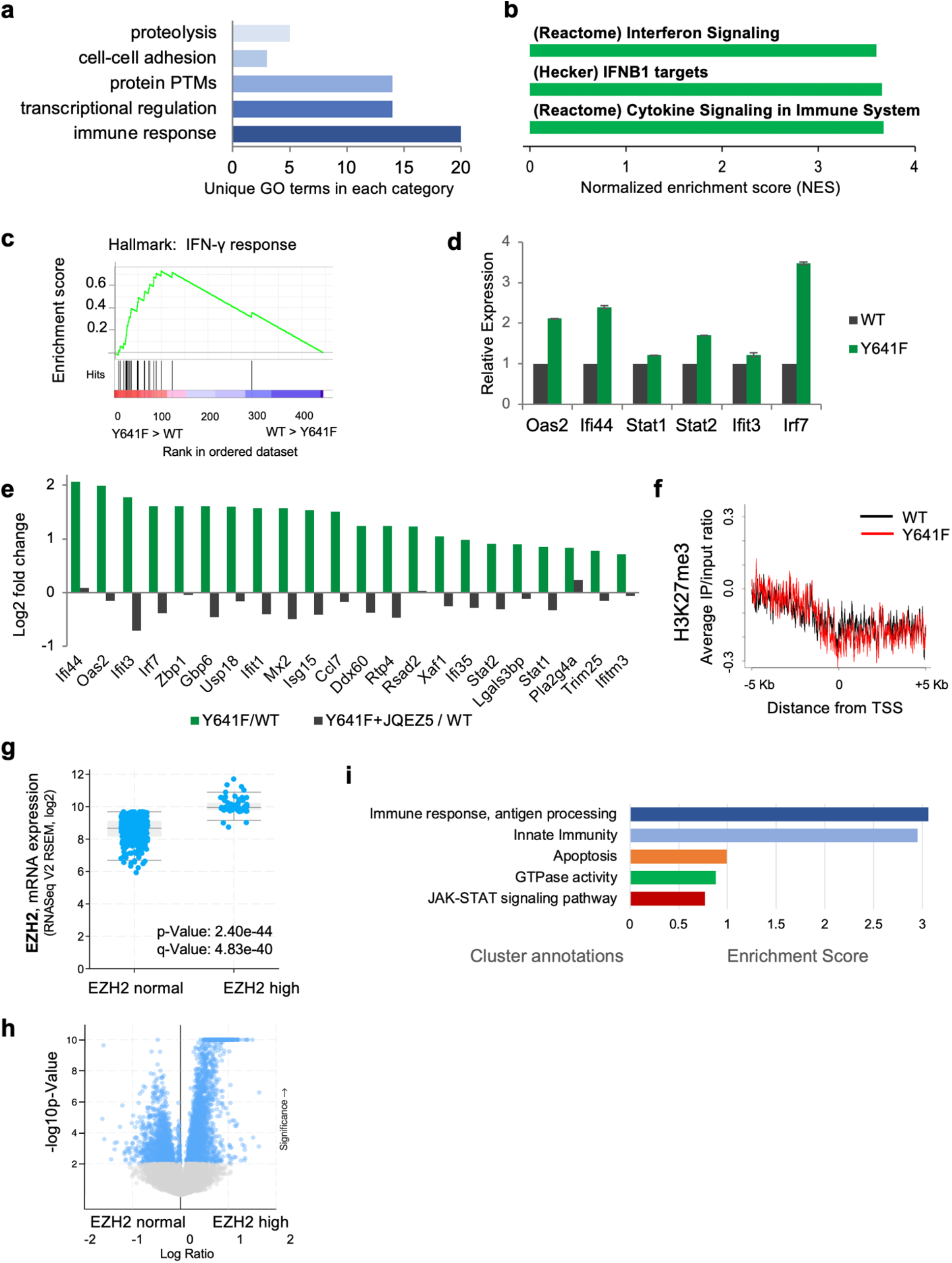
*Ezh2*^Y641F^ melanoma cells exhibit an interferon-related gene expression profile. **(a)** Functional annotation of gene expression profile analysis of *Ezh2*^WT^ vs *Ezh2*^Y641F^ mutant melanoma cells. Upregulated genes in *Ezh2*^Y641F^ cells were analyzed using the DAVID functional annotation clustering tool to identify enriched GO terms. **(b)** GSEA analysis of upregulated genes in *Ezh2*^Y641F^ melanoma cell lines showing enrichment for Interferon signaling categories. **(c)** Representative GSEA plot of the Hallmark: Interferon gamma response category. **(d)** Validation of differentially expressed interferon-related genes by qPCR **(e)** Log_2_ fold change of upregulated interferon-related genes in *Ezh2*^Y641F^ melanoma cells lines before and after treatment with Ezh2 inhibitor JQEZ5. **(f)** Analysis of H3K27me3 signals 5kb upstream and downstream of transcriptional start sites of the genes from (e) **(g)** EZH2 expression in human melanoma patients from TCGA split in two cohorts, EZH2-normal vs EZH2-high (includes exp>2 and *EZH2*^Y641^ mutations) **(h)** Volcano scatterplot of RNA-seq data from TCGA melanoma patients stratified by EZH2 expression (EZH2-normal vs EZH2-high). **(i)** Functional annotation analysis of downregulated genes in EZH2-high patients.

### Expression of IFN-related genes in Ezh2^Y641F^ melanomas is dependent on Ezh2 methyltransferase activity but not H3K27me3

Next, we wanted to determine if expression of these genes was dependent on Ezh2’s methyltransferase activity, so we analyzed expression of these genes after treatment with the Ezh2 inhibitor, JQEZ5. JQEZ5 is *S*-5′adenosyl-L-methionine (SAM)-competitive pyridinone inhibitor and is potent and selective against Ezh2 activity (13,36). We found that treatment with the Ezh2 inhibitor reversed the overexpression of these genes in the presence of the *Ezh2*^Y641F^ mutation, suggesting that their expression is dependent on Ezh2 activity (**Fig. 1e**). To determine whether expression of these genes was also dependent on the H3K27me3 mark that is mediated by enzymatically active Ezh2, we analyzed H3K27me3 distribution at the promoter regions of these differentially expressed genes. We expected that genes whose expression is increased in the *Ezh2*^Y641F^ mutants would exhibit a loss of the suppressive H3K27me3 mark along their promoters or gene bodies. Surprisingly, we did not find a decrease in H3K27me3, or any other correlation between expression of these genes and deposition of H3K27me3 at their promoters (**Fig. 1f**). These data suggest that even though expression of these IFN-related genes is dependent on Ezh2 activity, they are not dependent on its canonical function of methylating histone tails at H3K27, suggesting that expression of these genes is mediated through an indirect way, or a non-canonical function of Ezh2 activity.

### Immune response signatures in patients with hyperactive Ezh2

To determine whether the immune response pathways that we identified in mouse models reflect what happens in human patients, we analyzed gene expression data from TCGA melanoma studies. We identified 68/480 patients that expressed the *EZH2*^Y641^ mutant or exhibited increased EZH2 expression (exp>2) compared to the rest of the cohort (**Fig. 1g**). Functional annotation of the downregulated genes in the “EZH2-high” group (**Fig. 1h**) showed enrichment for immune response pathways, antigen recognition, apoptosis, GTPase activity, along with the JAK/STAT signaling pathway (**Fig. 1i**). These data are consistent with our mouse model data suggesting that our models are appropriate and physiologically relevant to further investigate the mechanistic details and consequences of these pathways in melanoma.

### Ezh2 directly interacts with Stat3 in melanoma cells

To understand how these IFN targets are upregulated in *Ezh2*^*Y641F*^ cells, we considered whether Ezh2 may interact with known IFN regulatory factors, such as the STAT proteins, especially since the JAK-STAT signaling pathways was identified in human melanoma patients with increase EZH2 activity (**Fig. 1g-i**). Additionally, previous studies have shown that Ezh2 directly interacts with and methylates Stat3 in several solid tumors (16,17,37), however, there was no consensus on the consequences of methylation on Stat3 activity. Yang et al. suggested that Stat3 methylation at lysine 140 (K140) inhibits Stat3 activity (37) whereas Dasgupta et al., reported that Ezh2-mediated methylation at lysine 49 (K49) was activating in colon carcinoma cell lines (16). In glioblastoma, Kim et al. reported that methylation at lysine 180 (K180) enhanced Stat3 activation (17). These differences could be explained by tissue-specific functionality of these proteins. To investigate the relevance of Stat3 in melanoma, we first determined whether Stat3 was expressed in our mouse model and melanoma cell lines and, indeed, Stat3 is present in both the cytoplasm and the nucleus (**Fig. 2a**). Interestingly, there was more Stat3 protein in the nucleus of *Ezh2*^*Y641F*^ cells than *Ezh2*^*WT*^, however, there was no difference in phosphorylated Stat3, suggesting no effect on canonical Stat3 activation (**Fig. 2a**). To test whether Ezh2 interacts with Stat3 in melanoma cells, we performed co-affinity immunoprecipitation in *Ezh2*^*WT*^ and *Ezh2*^*Y641F*^ cells using a Stat3 antibody and successfully pulled down both Ezh2 and Stat3 (**Fig. 2b**). This result was repeated with the Ezh2 antibody in all cell lines tested (**Fig. 2b**). Interestingly, when probing with a tri-methyl-recognizing antibody, we found that Stat3 was hypermethylated in the presence of Ezh2^Y641F^ protein. A much weaker interaction was observed in the presence of Ezh2^WT^ protein as well. These data suggest that Ezh2 directly interacts with Stat3, and is coincident with increased Stat3 methylation, which could contribute to the abnormal expression of IFN-related genes in *Ezh2*^*Y641F*^ melanomas shown in Figure 1.

**Figure 2.**
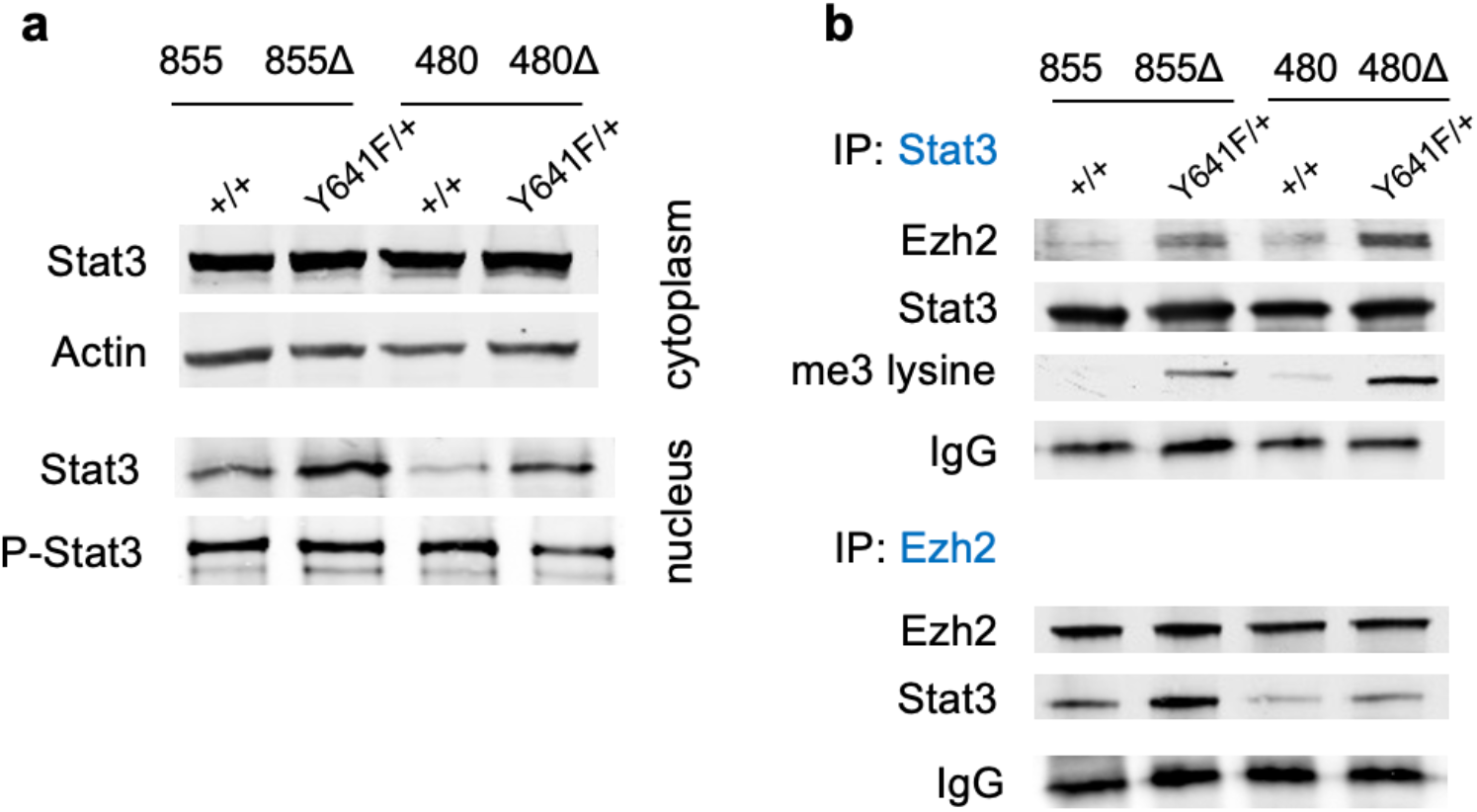
Stat3 physically interacts with Ezh2 in *Ezh2*^Y641F^ mutant cells and is hypermethylated. **(a)** Immunoblotting analysis for Stat3 and actin in cytoplasm, and Stat3 and phospho-Stat3 in nucleus of melanoma cell lines 480 and 855 (Δ = Y641F mutant). **(b)** Immunoprecipitation for Stat3 and Ezh2, immunoblotted for Stat3, Ezh2, and tri-methylated lysine.

### Stat3 knockdown in *Ezh2*^Y641F^ melanoma cells restores expression of IFN-regulated genes

To investigate the potential role of Stat3 in the differential gene expression patterns we observed in *Ezh2*^*Y641F*^ melanoma cells, we used lentiviral short hairpin RNA (shRNA) to knockdown Stat3 in the same melanoma cell lines used above (Fig. 3a). shRNA knockdown resulted in at least 90% knockdown efficiency and coincided with decreased expression of many of the upregulated IFN-related genes in Ezh2^Y641F^ cells (Fig. 3b), suggesting that the changes in IFN-related gene expression are at least partially mediated by Stat3.

**Figure 3.**
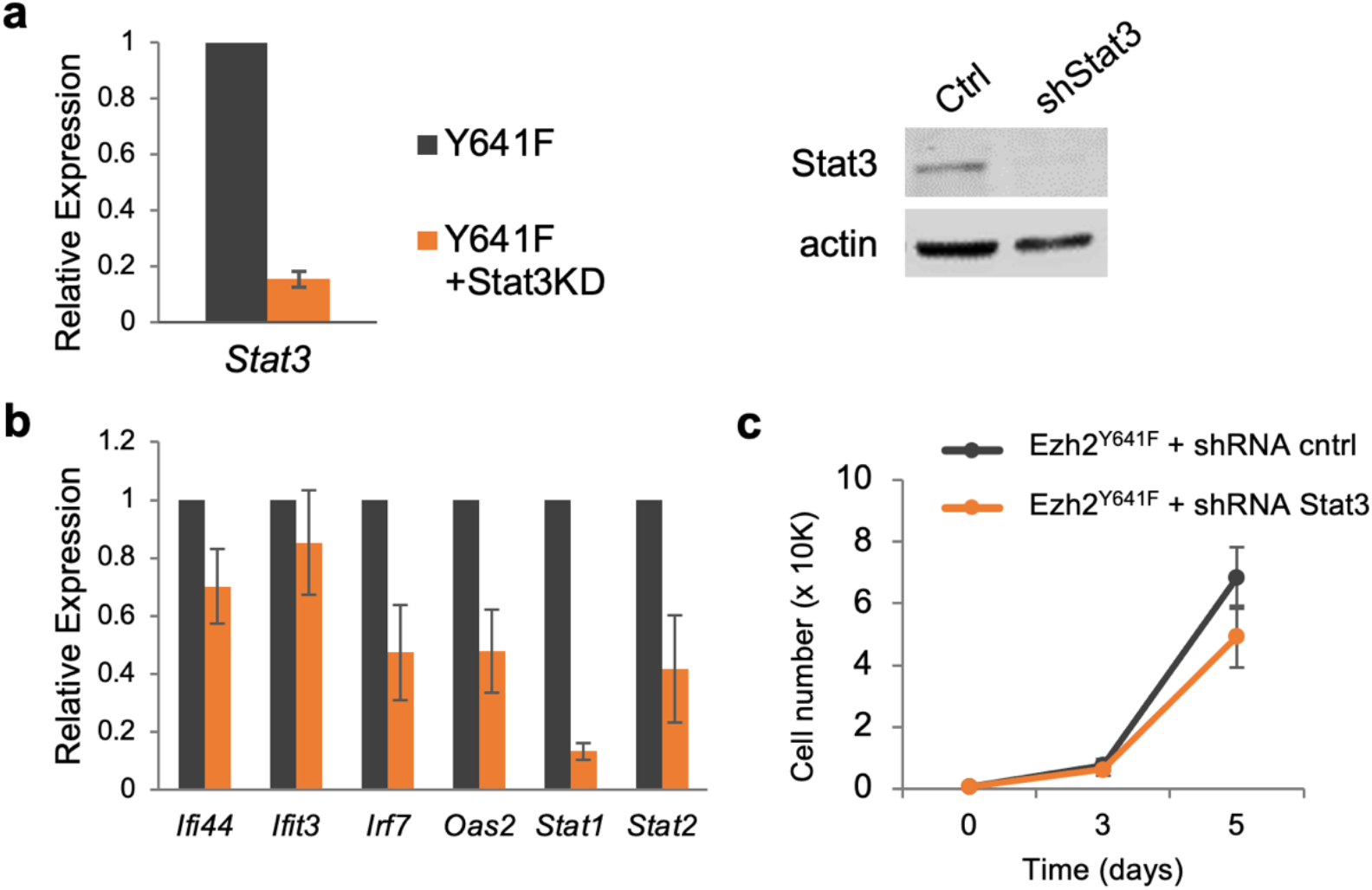
Stat3 knockdown in melanoma cell lines restores IFN gene expression without affecting intrinsic cell growth. **(a)** Lentiviral Stat3 knockdown (KD) in melanoma cell lines confirmed by Western blot and qPCR **(b)** Expression of interferon-related genes in control vs shRNA Stat3 knockdown. **(c)** Effect of shRNA Stat3 knockdown in *Ezh2*^Y641F^ melanoma cell lines on cell growth.

### Stat3 knockdown has no effect on *Ezh2*^*Y641F*^ melanoma cell-intrinsic growth

Stat3 can regulate a plethora of biological functions, including many cell-intrinsic hallmarks of cancer, such as cell growth, proliferation, apoptosis, and migration. To determine whether Stat3 regulates similar functions in melanoma, we performed growth assays in *Ezh2*^*Y641F*^ melanoma cell lines, with and without Stat3 knockdown. We found that Stat3 knockdown had no significant effect on *in vitro* cell growth of these melanoma cell lines (**Fig. 3c**), suggesting that if Stat3 plays a role in the pathogenesis of *Ezh2*^*Y641F*^ melanoma, it is not through regulation of intrinsic cell growth.

### Stat3 knockdown accelerates *in vivo* growth of *Ezh2*^*Y641F*^ melanoma tumors

An important aspect of tumor growth and progression is the interplay between the cancer cells and the tumor microenvironment. One of the many functions of Stat3 is regulation of inflammation, a cancer hallmark, which causes the microenvironment to be more supportive of tumor growth. Indeed, persistent Stat3 activation is a recurring feature in many epithelial tumors (26) and Stat3 expression and activation can affect the anti-tumor immune response. In most cancers, Stat3 activation and its downstream signaling has strong immunosuppressive properties (38), making Stat3 an attractive therapeutic target to augment the anti-tumor immune response.

To investigate the role of Stat3 in melanoma growth *in vivo*, we infected isogenic *Braf*^V600E^ *Pten*^F/+^ *Ezh2*^WT^ vs *Ezh2*^Y641F^ melanoma cell with anti-Stat3 shRNA or a scrambled control shRNA. These cell lines were transplanted orthotopically into recipient C57BL/6 mice and tumor growth was monitored over time. Knockdown of Stat3 in *Ezh2*^WT^ cell lines did not affect tumor growth, however, in the epigenetically altered background mediated by *Ezh2*^Y641F^, Stat3 knockdown led to increased tumor growth (**Fig. 4b**). Notably, this result in the *Ezh2*^Y641F^ mutant cell lines is the opposite of what one would expect based on the previously well-established immunosuppressive properties of activated Stat3 in solid tumors (26).

**Figure 4.**
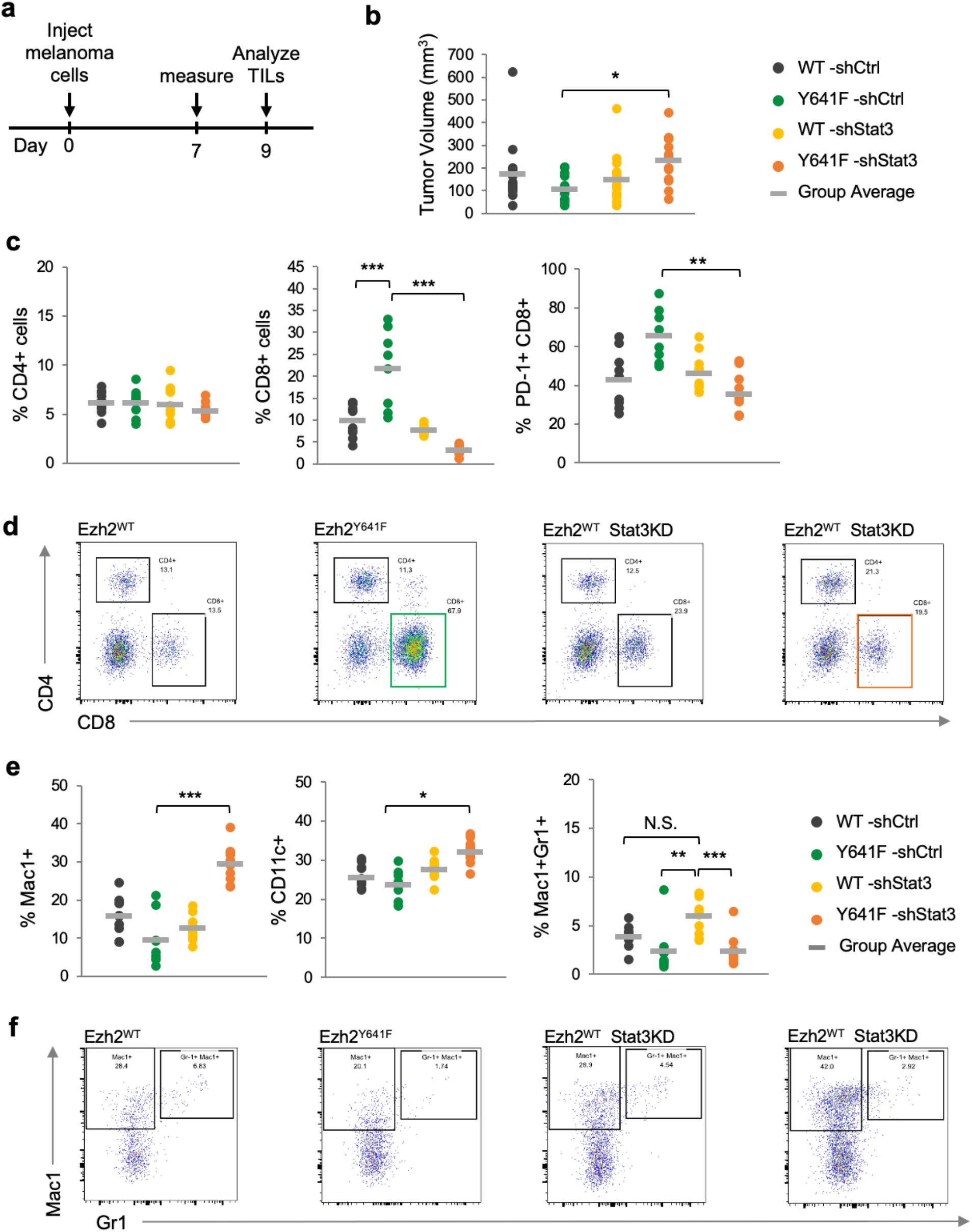
Analysis of tumor-infiltrating immune cells in *Ezh2*^WT^ vs *Ezh2*^Y641F^ melanoma cells after Stat3 knockdown. **(a)** Schematic of experimental design. Melanoma cells were injected subcutaneously into the flanks of C57Bl/6 mice. The tumors were measured 7 days later. N = 8-10/group. **(b)** *In vivo* tumor volume/growth over time of *Ezh2*^WT^vs *Ezh2*^Y641F^ melanoma cell lines, with and without Stat3 knockdown. **(c)** Analysis of tumor-infiltrated lymphocytes by flow cytometry, focusing on CD4+, CD8+ and CD8/PD-1+ cells. **(d)** Representative flow cytometry plots for panel (c). **(e)** Analysis of tumor-infiltrated myeloid cells by flow cytometry. **(f)** Representative flow cytometry plots for panel (e).

### Expression of *Ezh2*^Y641F^ in melanoma cells alters tumor-infiltrating immune populations which are dependent on continued Stat3 expression

To understand why *Ezh2*^Y641F^ mutant melanoma tumors were growing faster despite knocking down an immunosuppressive factor, we carefully analyzed the tumor microenvironment for the presence of immune cells. We dissected tumors from mice at day nine after initial injection and physically dissociated the tumors to isolate tumor-infiltrated hematopoietic cells for analysis via flow cytometry. Analysis for tumor-infiltrating lymphocytes revealed no differences in CD4+ cells, however, there was increased infiltration of cytotoxic CD8+ T cells in *Ezh2*^Y641F^ compared to *Ezh2*^WT^ tumors (**Fig. 4c**). Stat3 knockdown in *Ezh2*^WT^ tumors had no effect on the number of infiltrated CD8+ T cells. Stat3 knockdown, however, in *Ezh2*^Y641F^ tumors dramatically decreased the number of CD8+ cells in the tumor microenvironment (**Fig. 4c, d**), suggesting that expression of Stat3 becomes critical only when *Ezh2*^Y641F^ is expressed. Further analysis of these tumor-infiltrating T cells showed increased expression of the PD-1 receptor, suggesting that they may be exhausted. Expression of PD-1 on these cells was also dependent on expression of Stat3 (**Fig. 4c**).

In addition to lymphocyte populations, we also analyzed the tumor microenvironment for the presence of myeloid cells. There was an increase in CD11c+ dendritic cells and Mac1+ cells in *Ezh2*^Y641F^ cells after Stat3 knockdown, but not in *Ezh2*^WT^ cells. A different trend was observed with granulocytes (Mac1+ Gr1+ cells), where Stat3 knockdown caused a non-significant increase (compared to *Ezh2*^WT^ without knockdown) in the *Ezh2*^WT^ tumors (**Fig. 4e, f**). Taken together, these data suggest that expression of *Ezh2*^Y641F^ in these tumors has created a unique dependency on Stat3 expression, with significant consequences on the tumor microenvironment.

### Inhibition of Stat3 expression limits the anti-tumor immune response and promotes melanoma recurence

The changes in the composition of the tumor microenvironment described above suggest that Stat3 plays a direct role in the growth and progression of these melanoma tumors *in vivo*. To understand the role of the tumor microenvironment and infiltration of immune cells on tumor growth, we utilized the same *in vivo* tumor model described above and observed tumor growth over a longer period. Typically, these tumors grow rapidly for about ten days, but then most are spontaneously rejected by the host (**Fig. 5a**) and shrink to undetectable size by palpation by day 25. However, some tumors recur, at which point the host can no longer inhibit their growth and the animals must be euthanized. Interestingly, cell lines with Stat3 knockdown exhibit a much higher incidence of persistent or recurrent tumor growth (**Fig 5b, c**), regardless of *Ezh2* genotype. Pharmacologic inhibition of Stat3 using the small molecule STATTIC (39) had the same effect as genetic Stat3 knockdown (**Fig. 5d**), suggesting that the effects we observe are not off-target effects of the shRNA system. To determine if the tumor rejection was due to the presence of cytotoxic T cells in the tumor microenvironment, we depleted the CD8+ cells in tumor-bearing animals by administering an anti-CD8 antibody. Depletion of CD8+ cells prevented rejection of the tumors, which continued to grow rapidly compared to vehicle-treated controls (**Fig. 5e, f**). Additionally, we injected the same cell lines into immunodeficient NSG mice and monitored growth over time. In this setting, there was no rejection and tumors continued to grow rapidly (**Fig. 5g**), clearly suggesting that the effects on tumor growth dynamics that we observe are mediated by the immune system.

**Figure 5.**
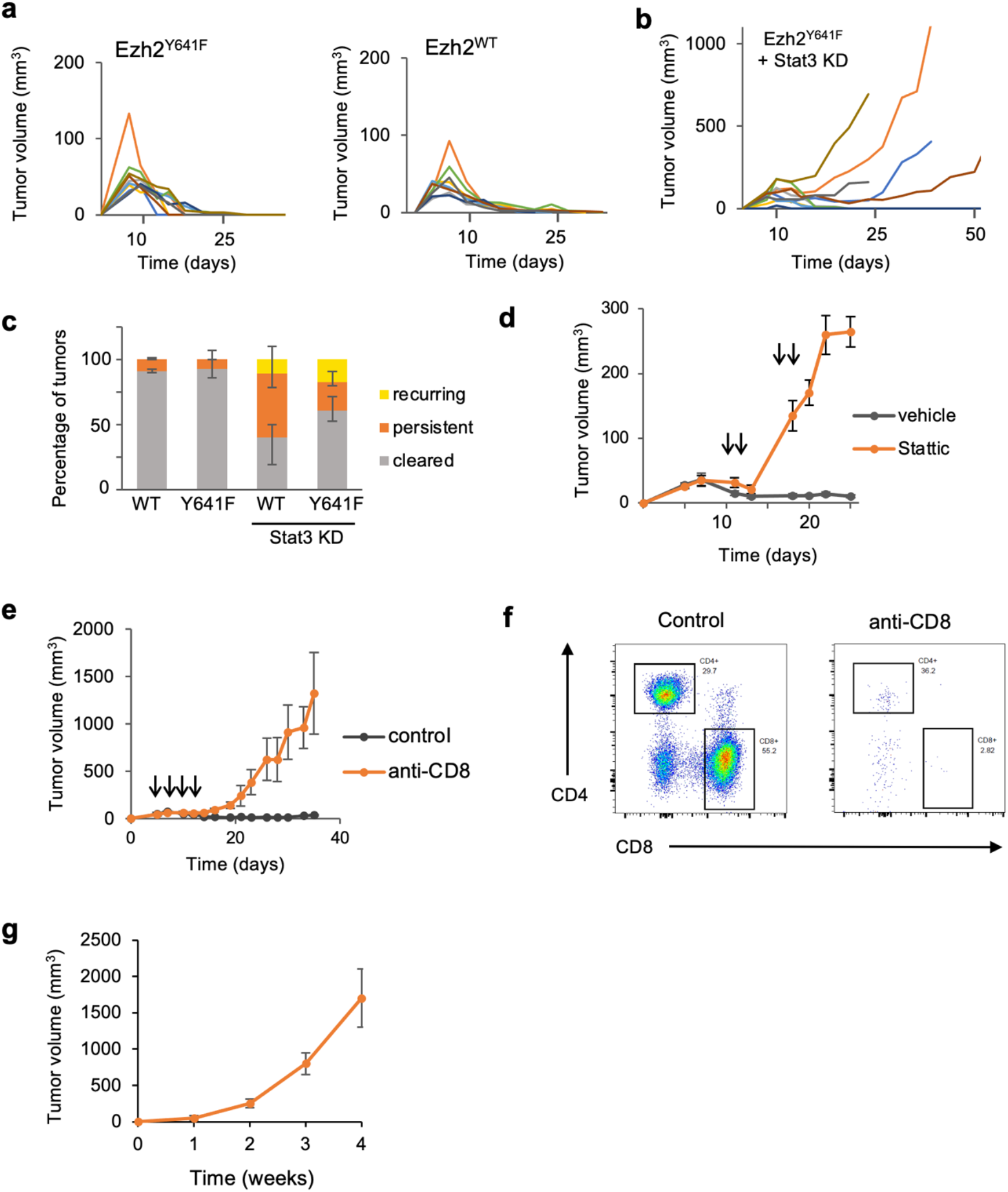
Melanoma tumor growth over time in *Ezh2*^WT^ vs *Ezh2*^Y641F^ melanoma cells after Stat3 knockdown. **(a)** Tumor volume over time for the indicated genotypes. Mice are on a *Braf*^V600E^ *Pten*^F/+^ background. Each line represents an individual tumor. N = 10/group. These graphs are from one of two replicate experiments. **(b)** Tumor volume over time for *Ezh2*^Y641F^ cell lines after Stat3 knockdown. N = 10/group. These graphs are from one of two replicate experiments. **(c)** Tumor outcome average of two replicate experiments, stratified into 3 groups: cleared (rejected tumors), persistent (tumors that grew consistently), and recurring (tumors that shrunk but then rebounded). N = 10-14/group. **(d)** Tumor growth over time after treatment with Stat3 inhibitor, STATTIC. **(e)** Tumor volume over time with anti-CD8 treatment. N = 8/group. Arrows indicate the time of anti-CD8 treatments. **(f)** Representative plots of CD4 and CD8 expression in tumor-infiltrating immune cells from tumors treated with or without anti-CD8 antibody. **(g)** Tumor volume from an *Ezh2*^Y641F^ melanoma cell line from (a), injected in NSG mice. N = 5.

### *Ezh2*^Y641F^ changes global Ezh2 and Stat3 association with chromatin and DNA

To understand the molecular effects and the nature of the interaction between Ezh2 and Stat3 in melanoma, we performed chromatin immunoprecipitation followed by sequencing (ChIP-seq) for both Ezh2 and Stat3 in *Ezh2*^WT^ vs *Ezh2*^Y641F^ melanoma cells. Three separate syngeneic cell lines were used for these experiments as biological controls. Comparative analysis between genotypes revealed several global differences mediated by expression of *Ezh2*^Y641F^. For example, in *Ezh2*^Y641F^ melanoma cells, Ezh2 protein was bound less frequently on promoter regions but more within the first intron (**Supp. Fig. 2a, c**). These data are consistent with our previous observations where H3K27me3 in *Ezh2*^Y641F^ is redistributed away from promoters and towards gene bodies (13). Stat3 binding across the genome was also altered in *Ezh2*^Y641F^ with less binding in promoter regions compared to *Ezh2*^WT^ (**Supp. Fig. 2b, d**). Systematic analysis of all differential Stat3 peaks showed binding/association of Stat3 with new genomic loci in *Ezh2*^Y641F^ cells (**Supp. Fig. 2e**). Analysis of called peaks within proximal genes in Ezh2 and Stat3 ChIP identified numerous already known loci that are bound by either protein. For example, Ezh2 was associated with known PRC2-regulated regions, such as *Hoxa* and *Hoxc* clusters, and Stat3 was associated with known Stat3 targets, such as *Hif1a, Pim2, Foxo1, Il10* (40) (**Supp. Fig. 3**).

Next, we investigated the functional significance of the binding sites that were identified by ChIP-seq. Very often, associating only proximal binding events (2-5 kb from the transcriptional start site), results in discarding a significant fraction of the binding events (30), and associating each binding site with the nearest gene also introduces biases and many false positives. Therefore, we used the Genomic Regions Enrichment of Annotations Tool (GREAT) to analyze the functional significance of cis-regulatory regions with Ezh2 and Stat3 binding by modeling the vertebrate genome regulatory landscape from multiple sources (30). Using the GREAT algorithm, we obtained a list of genes previously shown to be regulated by regions bound by Ezh2 and Stat3. Gene set enrichment analysis of Ezh2 peaks showed enrichment for known and previously identified PRC2 and EZH2 targets (**Supp. Fig. 4a**), supporting the quality of the ChIP-seq data. Functional annotation analysis identified multiple pathways implicated in cancer cell-intrinsic and cell-extrinsic mechanisms, such as endothelial cell migration, leukocyte differentiation and regulation of IL-6 signaling, along with other pathways not obviously implicated in oncogenic mechanisms (**Supp. Fig. 4b-d**).

### Ezh2 and Stat3 bind to common regions across the genome

To understand if Ezh2 and Stat3 co-operate in regulating gene expression, we investigated whether Ezh2 and Stat3 bind to overlapping genomic regions. Comparison of ChIP-seq peaks for Ezh2 and Stat3 in *Ezh2*^Y641F^ cells identified unique loci co-occupied by Ezh2 and Stat3 (**Fig. 6a**). Binding to these loci was not detected in *Ezh2*^WT^ cells. Motif analysis of these binding sites revealed links to melanoma initiation pathways, such as Mitf binding motifs, in addition to immune regulatory elements, such as Stat4 and Irf6 **(Fig. 6b**), consistent with the role for Ezh2 and Stat3 in regulating the tumor immune response in these cells.

**Figure 6.**
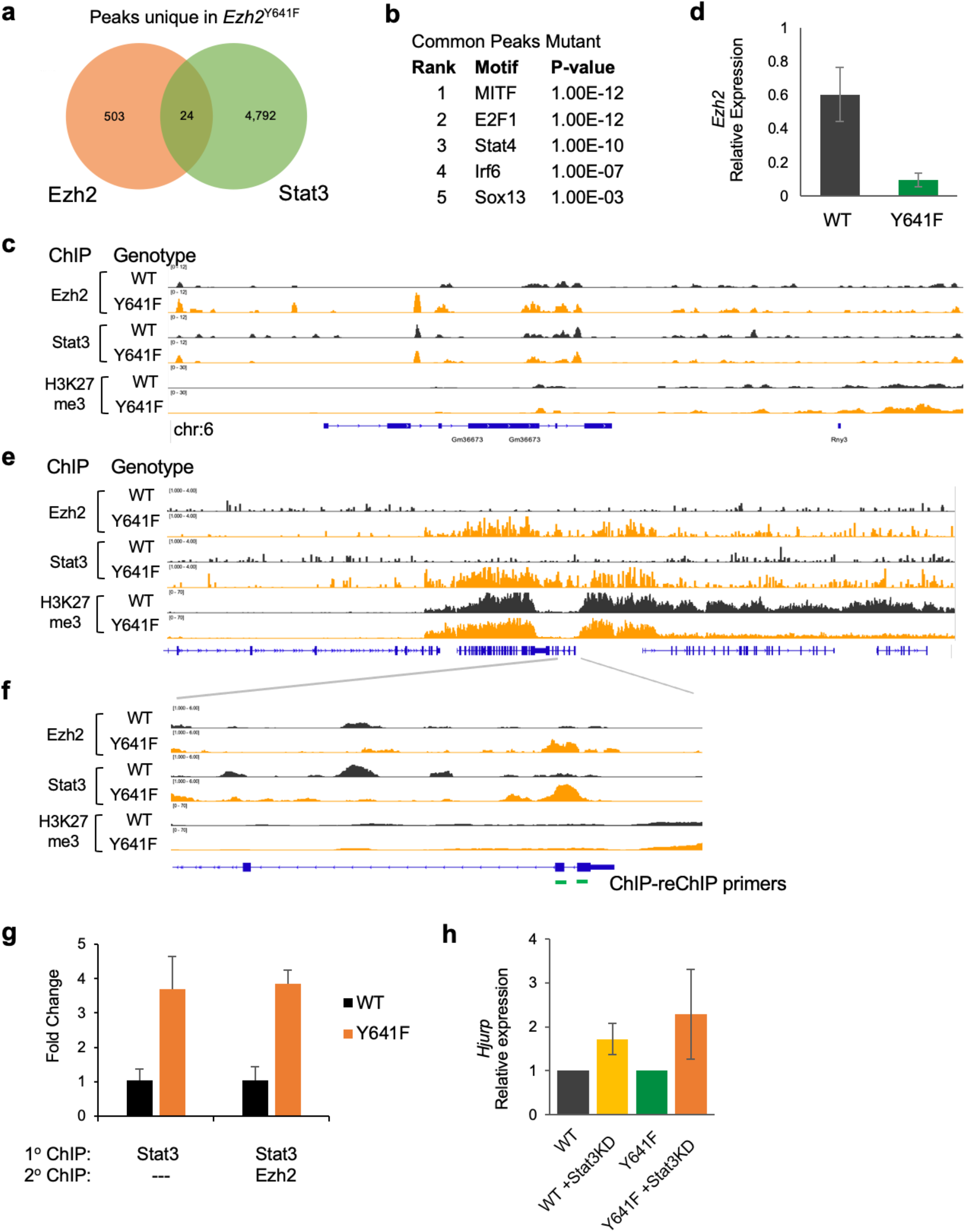
Changes in global DNA and chromatin binding by Ezh2 and Stat3 in *Ezh2*^Y641F^mutant melanoma cells. **(a)** Distinct and overlapping ChIP-seq peaks for Ezh2 (left) and Stat3 (right) in *Ezh2*^Y641F^melanoma cells. **(b)** Motifs identified in the common peaks between Ezh2 and Stat3 in *Ezh2*^Y641F^ melanoma cells from panel (a). **(c)** Ezh2, Stat3 and H3K27me3 ChIP-seq peaks at a regulatory region on chromosome 6, 72kb upstream of *Ezh2* that regulate its expression. **(d)** Expression of *Ezh2* in *Ezh2*^WT^ vs *Ezh2*^Y641F^ melanoma cells after Stat3 knockdown. **(e)** ChIP-seq peaks of Ezh2 and Stat3 in *Ezh2*^Y641F^ cells on chromosome 1, proximal to genes *Hjurp* and *Trpm8*. H3K27me3 ChIP-seq tracks also visualized for comparison. **(f)** Zoomed-in region of the *Hjurp* promoter highlighting new Ezh2 and Stat3 binding sites at the *Hjurp* promoter in *Ezh2*^Y641F^ melanoma cells. **(g)** ChIP-qPCR and ChIP-reChIP assay at the *Hjurp* promoter, with anti-Stat3 and anti-Ezh2 antibodies. **(h)** *Hjurp* gene expression in *Ezh2*^WT^ and *Ezh2*^Y641F^ melanoma cells after Stat3 knockdown (KD). Expression of knockdown groups is shown relative to the controls.

### *Ezh2* expression becomes dependent on the presence Stat3 in *Ezh2*^Y641F^ melanoma cells

Interestingly, one locus co-occupied by Ezh2 and Stat3 was on chromosome 6, 70 kb upstream of the genomic locus of the *Ezh2*, and was predicted to regulate expression of *Ezh2* itself (30). Even though Ezh2 protein binds to that region, no significant H3K27me3 mark was associated with it (**Fig 6c**), suggesting that expression of *Ezh2* in *Ezh2*^Y641F^ cells might be dependent on the interaction between Ezh2 and Stat3. To determine whether *Ezh2* expression was dependent on the presence of Stat3, we knocked down Stat3 in multiple melanoma cell lines and assayed *Ezh2* expression. Consistent with the ChIP-seq data and the long-range interaction predicted by the GREAT algorithm, Stat3 knockdown caused a ten-fold decrease in *Ezh2* expression in *Ezh2*^Y641F^ cells, while only mildly decreasing expression in *Ezh2*^WT^ cells (**Fig. 6d**), further highlighting this unique interaction between Ezh2 and Stat3 in *Ezh2*^Y641F^ melanoma cells.

### Ezh2 and Stat3 bind at common genomic loci and repress gene expression

Binding of both Ezh2 and Stat3 at the co-occupied loci proximal to gene start sites and within canonical promoter regions might result in direct regulation of transcription. We thus directly investigated several of these loci with ChIP-qPCR. An example is shown in Figure 6d and includes the genes *Hjurp* and *Trpm8*. To verify that Ezh2 and Stat3 bind to that region, we performed ChIP followed by qPCR using primers that corresponded to the coordinates of specific peaks. Additionally, to determine if Ezh2 and Stat3 bind together at the above regions, we performed sequential ChIP (ChIP-reChIP), providing further evidence that Ezh2 and Stat3 co-localize at these genomic loci (**Fig. 6g**). To determine the effect of Ezh2 and Stat3 binding on the expression of the genes most proximal to ChIP-seq peaks, we performed qPCR in *Ezh2*^WT^ vs *Ezh2*^Y641F^ cells with and without shRNA-mediated Stat3 knockdown. We found that *Trpm8* was not expressed in either genotype, and its expression did not change after Stat3 knockdown. *Hjurp* expression was readily detectable, and was decreased in *Ezh2*^Y641F^ cells. Upon Stat3 knockdown, *Hjurp* expression significantly increased in both *Ezh2*^WT^ and *Ezh2*^Y641F^ cells, but more significantly in *Ezh2*^Y641F^ cells (**Fig. 6h**). These data suggest that the presence of both Ezh2 and Stat3 at these loci suppresses *Hjurp* expression, likely by recruiting the entire PCR2 complex as this region is also heavily marked by H3K27me3 (**Fig. 6e**).

### Stat3 interacts with Ezh2 within the context of the rest of the PRC2 complex

Previous studies have demonstrated that Stat3 binding to its target genes is not always activating in nature but also repressive (24). The suppressive properties of Stat3 could be dependent on association other proteins, co-factors or complexes, however, no such mechanistic insight has previously been shown. The results in figure 6e-g indicate that Stat3 association at the *Hjurp* locus is repressive, and the presence of abundant H3K27me3 suggests that suppression of *Hjurp* expression might be mediated via association with the repressive chromatin-modifying complex, PRC2. To determine whether the Stat3 interaction with Ezh2 takes place within the context of the entire PRC2 complex, we performed co-affinity immunoprecipitation assays using a Stat3 antibody and assayed for interactions with Suz12 and Eed, the other two core components of the PRC2 complex. We found that Stat3 interacts with both Suz12 and Eed in both *Ezh2*^WT^ and *Ezh2*^Y641F^ cells, but the interaction with the mutant *Ezh2*^Y641F^ was stronger (**Fig. 7a, b**), suggesting that a possible mechanism of how Stat3 represses gene expression may be through association with the PRC2 complex.

**Figure 7.**
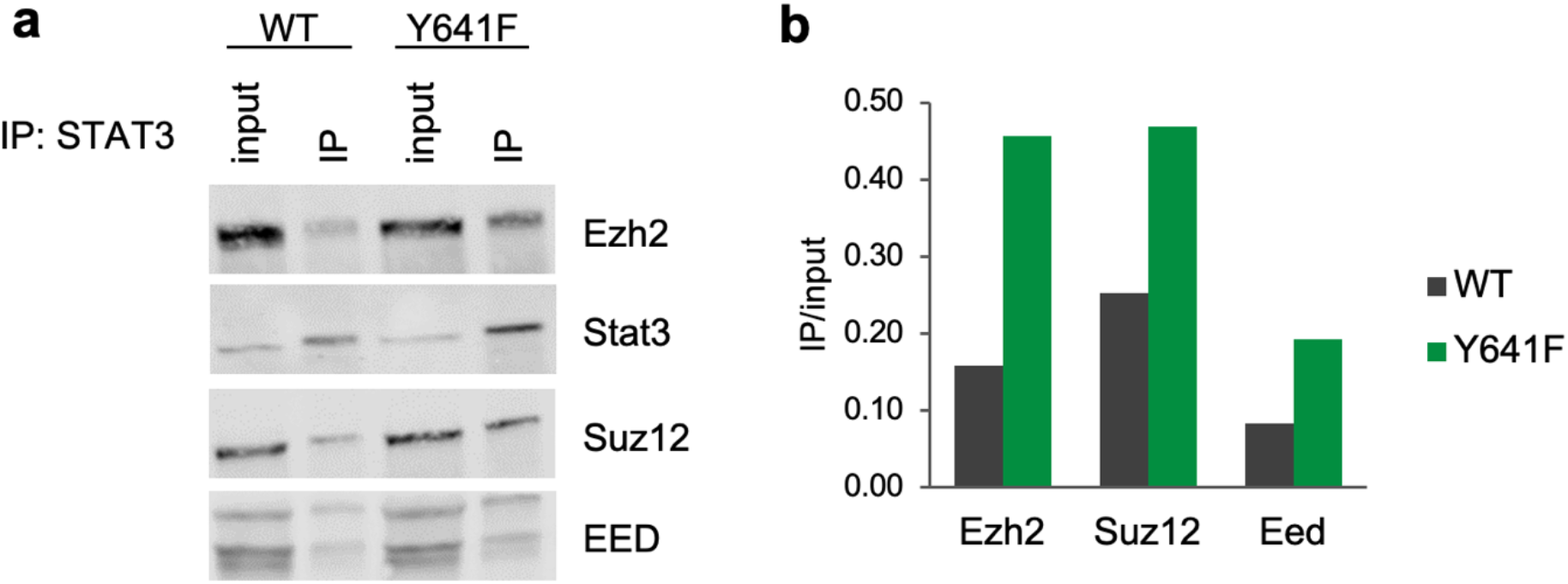
Stat3 protein interactions with core components of the PRC2 complex. **(a)** Immunoblotting analysis of Ezh2, Stat3, Suz12 and Eed after immunoprecipitation with anti-Stat3 antibody in *Ezh2*^WT^ vs *Ezh2*^Y641F^ melanoma cell lines. **(b)** Quantitative analysis of the blots in (a).

## DISCUSSION

To understand the oncogenic mechanisms of *Ezh2* in melanoma, we investigated the role of hyperactive *Ezh2* mutations in the anti-tumor immune response and also examined possible non-canonical interactions. Here we report that *Ezh2*^Y641F^ mutations promote a non-canonical interaction between Ezh2 and Stat3, with significant consequences on the anti-tumor immune response in melanoma. Specifically, expression of Stat3 is required for *Ezh2*^Y641F^-mediated infiltration of lymphocytes in the tumor microenvironment and consequently, for maintaining an anti-tumor immune response. Molecularly, we show that expression of mutant *Ezh2*^Y641F^ changes recognition and binding of Ezh2 and Stat3 across the genome and that the two proteins share common binding loci with complicated effects on gene expression. Together, these data suggest that the tumor microenvironment is a significant factor in the oncogenic properties of Ezh2 in melanoma.

Even though numerous sequencing studies have identified frequent mutations in epigenetic and chromatin-modifying factors in many cancers, their downstream oncogenic mechanisms remain underexplored. Understanding these mechanisms is critical in identifying which patients might respond to treatment, which drug combinations would be beneficial, and whether any synthetic lethal interactions exist that could be manipulated therapeutically. Many small molecule inhibitors targeting epigenetic modifiers are currently approved for use in patients, a number of which have significant benefits. Many patients, however, do not respond to treatment. The same is true for immunotherapy. Even though immunotherapy approaches have had great success in many melanoma patients, a significant portion of patients do not respond to treatment or exhibit significant side effects that prevent further use of these compounds. Understanding these mechanisms is therefore very important in developing therapeutic strategies for these patients.

Prior studies have implicated the role of Ezh2 in the anti-tumor immune response (15). For example, Zingg et al. demonstrated a high correlation between tumor T cell infiltration and high EZH2-PRC2 complex activity in human skin cutaneous melanoma. They proposed that Ezh2 silenced the tumors’ own immunogenicity and antigen presentation, contributing towards melanoma progression. In this study, we show that *Ezh2*^Y641F^ mutant melanomas also exhibit increased infiltration of cytotoxic T cells, whose presence in the melanoma microenvironment is dependent on the expression of the hyperactive *Ezh2*^Y641F^ mutant and on continued expression of Stat3. There are, therefore, multiple pathways through which Ezh2 may be modulating the anti-tumor immune response in melanoma, and these may also be dependent on the rest of the genetic/epigenetic makeup and specific oncogenic drivers expressed in melanoma or the models used in experiments. Since in human melanoma patients *Ezh2*^Y641F^ mutations almost always co-occur with *Braf*^V600E^ mutations, we combined conditional *Ezh2*^Y641F^ and *Braf*^V600E^ mouse alleles to generate the most physiologically relevant model. Mouse genetic studies corroborated our observations in human patients, where the *EZH2*^Y641F^ mutations co-operate with *BRAF* mutations but not *NRAS* mutations to accelerate the onset of melanoma (13). On the other hand, in the Zingg et al. studies, an *Nras*-driven melanoma model was used to study immune responses related to Ezh2 expression. Different results between the studies could be dependent on secondary oncogenic drivers and the genetic background of the cell lines used. Despite these differences, Ezh2 is undeniably implicated in the anti-tumor immune response, possibly utilizing more than one pathway.

The direct physical interaction between Ezh2 and Stat3 has previously been observed in other solid tumor models, however, the functional significance of that interaction has remained unexplored (16,17,37). Additionally, there was no consensus on whether Stat3 methylation promotes or inhibits phosphorylation and activation of Stat3. In the melanoma mouse models used in this study, Stat3 methylation does not influence Stat3 phosphorylation, and consequently its activation. This is further supported by the fact that gene expression analysis did not reveal a specific Stat3 signature, suggesting that Ezh2-mediated methylation of Stat3 does not have a significant effect on canonical Stat3 activity. Our data suggest that Stat3 physically interacts with Ezh2 and other PRC2 core components and is associated at common and distinct loci. Furthermore, we show that some loci where Ezh2 and Stat3 interact are also marked with H3K27me3, the repressive chromatin mark deposited by PRC2. At the same time, there are loci where Ezh2-Stat3 binding is not associated with the presence of H3K27me3, such as a distant regulatory region that controls expression of *Ezh2* itself via long-range chromatin interactions. It is possible, therefore, that the Ezh2-Stat3 interaction has a dual nature. When interacting with the entire PRC2 complex, Stat3 is associated with regions that will be marked with H3K27me3 and are thus repressed. This is consistent with the fact that Stat3 can also repress many of its target genes (24), although the mechanisms of target gene repression has so far remained unknown. On the other hand, Ezh2 and Stat3 could form a dimer without association with the PRC2 complex and activate transcription. This is also supported by data from prostate cancer studies, where the oncogenic function of Ezh2 is independent of the PRC2 complex and it involves the ability of Ezh2 to act as a coactivator with an intact methyltransferase domain to regulate expression of the androgen receptor (41).

The interaction of Stat3 with epigenetic regulators has been demonstrated in multiple studies. As stated above, and in this study, Stat3 can interact with components of the PRC2 complex. Previous studies in embryonic stem cells have shown that Stat3 is dependent on Brg1 (Smarca4), the ATPase subunit of the chromatin remodeling complex BAF (SWI/SNF) to activate the pluripotent genome (42,43). The SWI/SNF complex typically antagonizes PRC2, however, a recent study proposed that it can also facilitate PRC2 activity (44). Since Stat3 appears to interact with both complexes, it is possible that Stat3 is an important aspect of the relationship between the two chromatin-modifying complexes.

In summary, we show that expression of the hyperactive Ezh2^Y641F^ mutant in melanoma cells has a direct effect on the melanoma anti-tumor immune response and is dependent on expression of Stat3. This study also highlights the non-canonical functions of hyperactive Ezh2. However, several questions remain unanswered. For example, we do not understand the biochemical details of how Ezh2 may be directly methylating Stat3, which lysine is methylated, whether it is it mono-, di- or tri-methylated, or whether Stat3 methylation requires the entire PRC2 complex. We also do not know if Stat3 methylation is in fact consequential, and not merely a transient interaction enhanced by the presence of a hyperactive Ezh2 protein. Despite these unanswered questions, the interaction between Ezh2 and Stat3 will clearly have clinical importance. Stat3, which typically suppresses the anti-tumor immune response in other solid tumors, has the opposite effect in *Braf*^V600E^ Pten^F/+^*Ezh2*^Y641F^ melanomas. The cellular, genetic, and epigenetic context must therefore be taken into consideration when assessing the possible effect of manipulating these pathways for therapeutic purposes. More studies are needed to thoroughly investigate the extent to which the immunomodulatory properties of Stat3 may depend on other genetic or epigenetic pathways. This work, therefore, opens new avenues of investigation in understanding the biochemical, molecular, and clinical implications of these interactions and will have a significant impact on patient stratification and in the design of future therapeutic strategies.

## ACKNOWLEDGEMENTS

We thank the Siteman Flow Cytometry facility, McDonnel Genome Institute/Genome Access Center for technical assistance, and the Department of Comparative Medicine for animal expertise. We also thank all members of the Souroullas lab for critical input on the manuscript. This work was supported by the US National Cancer Institute K22-CA229612-01(GPS), T32 CA113275-10 (SZ), and the Cancer Research Foundation, Chicago IL, (GPS).

## AUTHOR CONTRIBUTIONS

GPS and SMZ designed experiments and wrote the manuscript. GPS, SJN and SMZ performed experiments, analyzed, and interpreted the data. LR, PYC, RLP, SS and PNL performed experiments. GPS conceived of and supervised the study.

## CONFLIC OF INTEREST

The authors declare no relevant competing financial interests.

## SUPPLEMENTARY FIGURES AND LEGENDS

**Supp. Fig 1.**
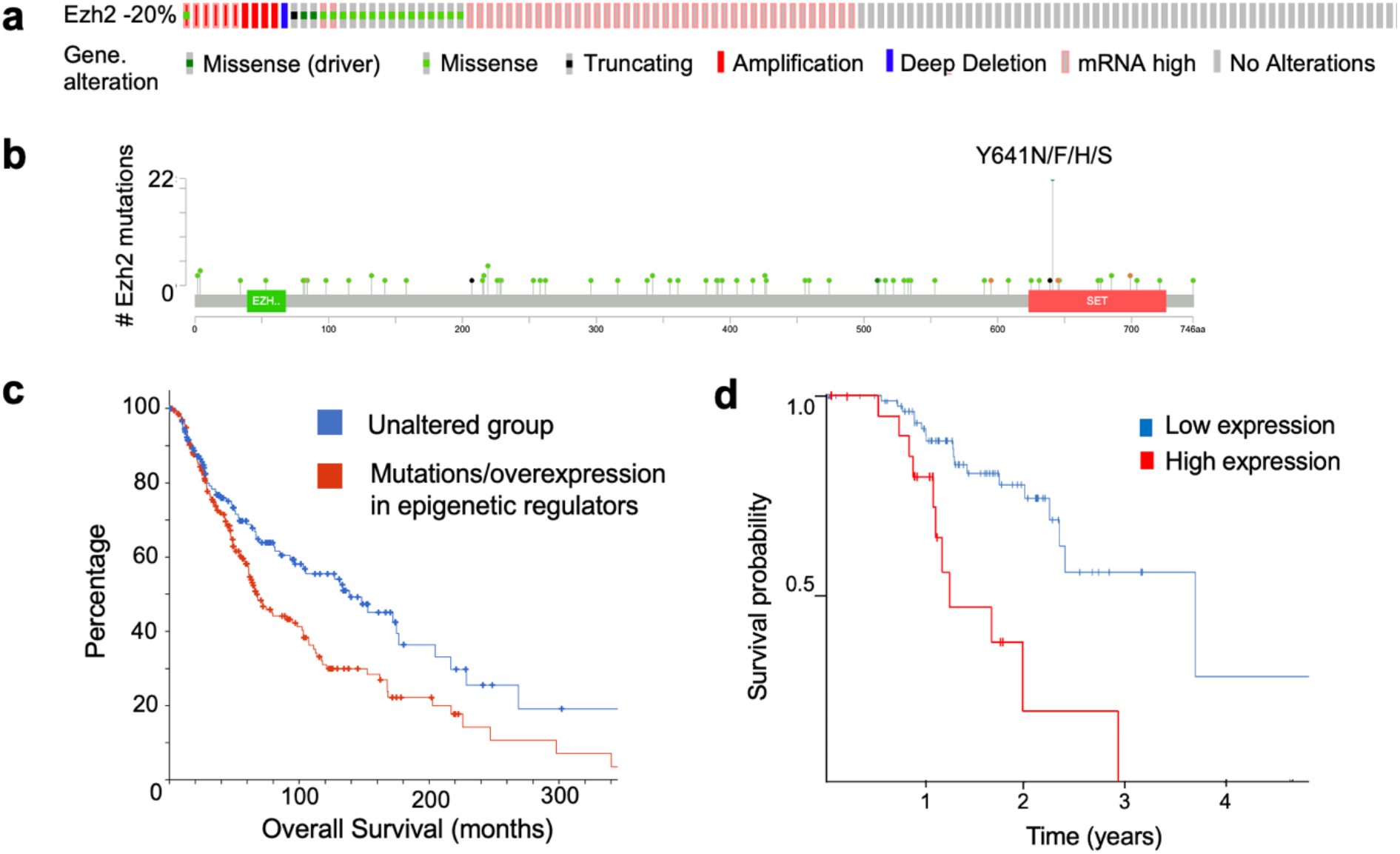
EZH2 expression and overall survival in human melanoma. **(a)** Frequency of EZH2 overexpression (exp>2) and activating Ezh2^Y641F^ mutations in human melanoma (data from TCGA) **(b)** Distribution of point mutations in EZH2 from TCGA data. **(c)** Overall survival of melanoma patients with PRC2 mutations/overexpression and other closely related complexes (SWI/SNF, NSD, SETD2) **(d)** Overall survival in human melanoma patients with higher-than-average EZH2 protein expression (data from Protein Atlas).

**Supp. Fig 2.**
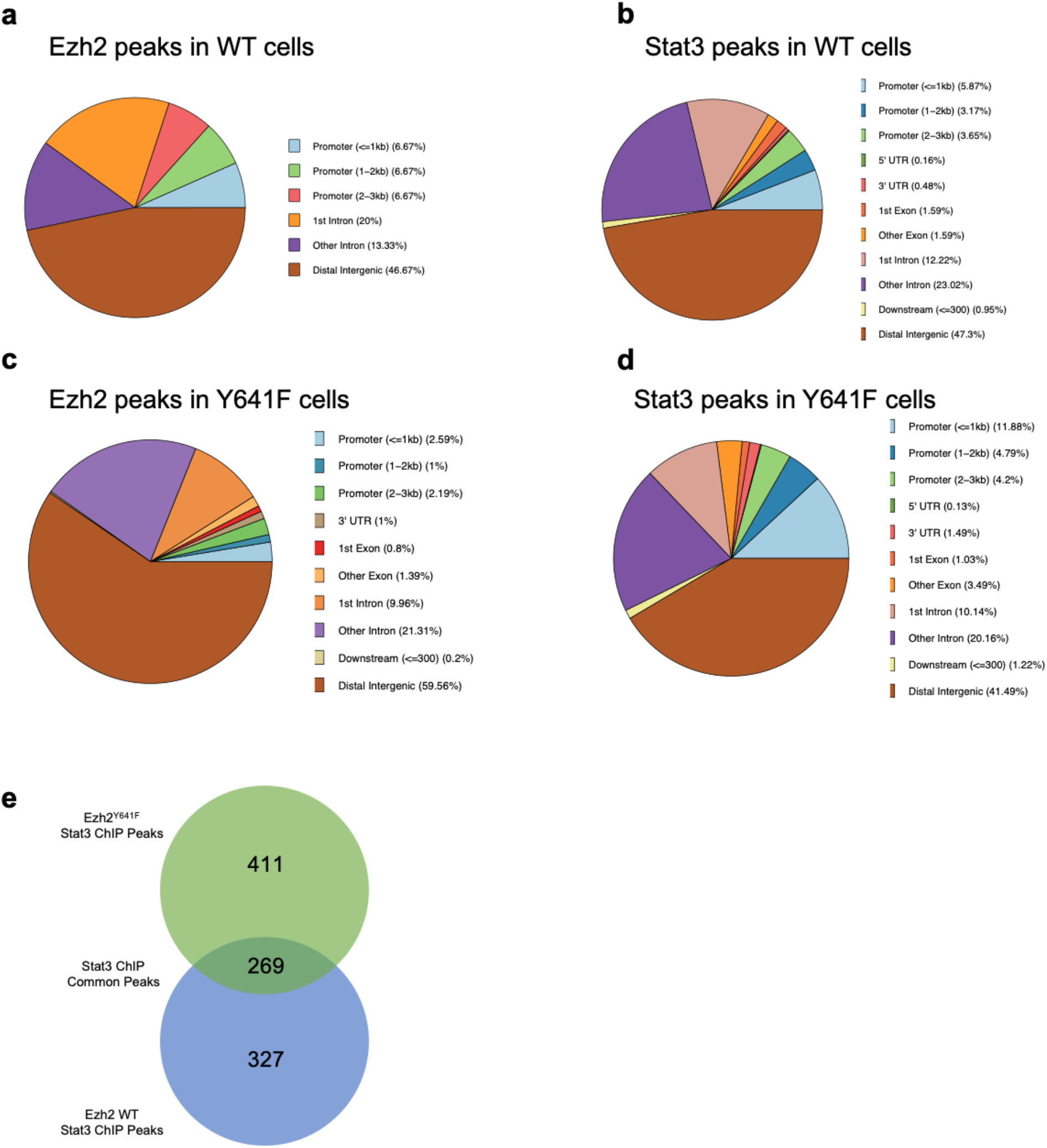
Global patterns of peak distribution in Ezh2 and Stat3 ChIP-seq data. Distribution of ChIP-seq peaks with the indicated genotypes: **(a)** Ezh2 peaks in Ezh2^WT^ cells, **(b)** Stat3 peaks in WT cells, **(c)** Ezh2 peaks in Ezh2^Y641F^ cells, **(d)** Stat3 peaks in Ezh2^Y641F^ cells **(e)** Stat3 ChIP-seq peaks in Ezh2^Y641F^ (top-green) vs Ezh2^WT^ (bottom-blue)

**Supp. Fig 3.**
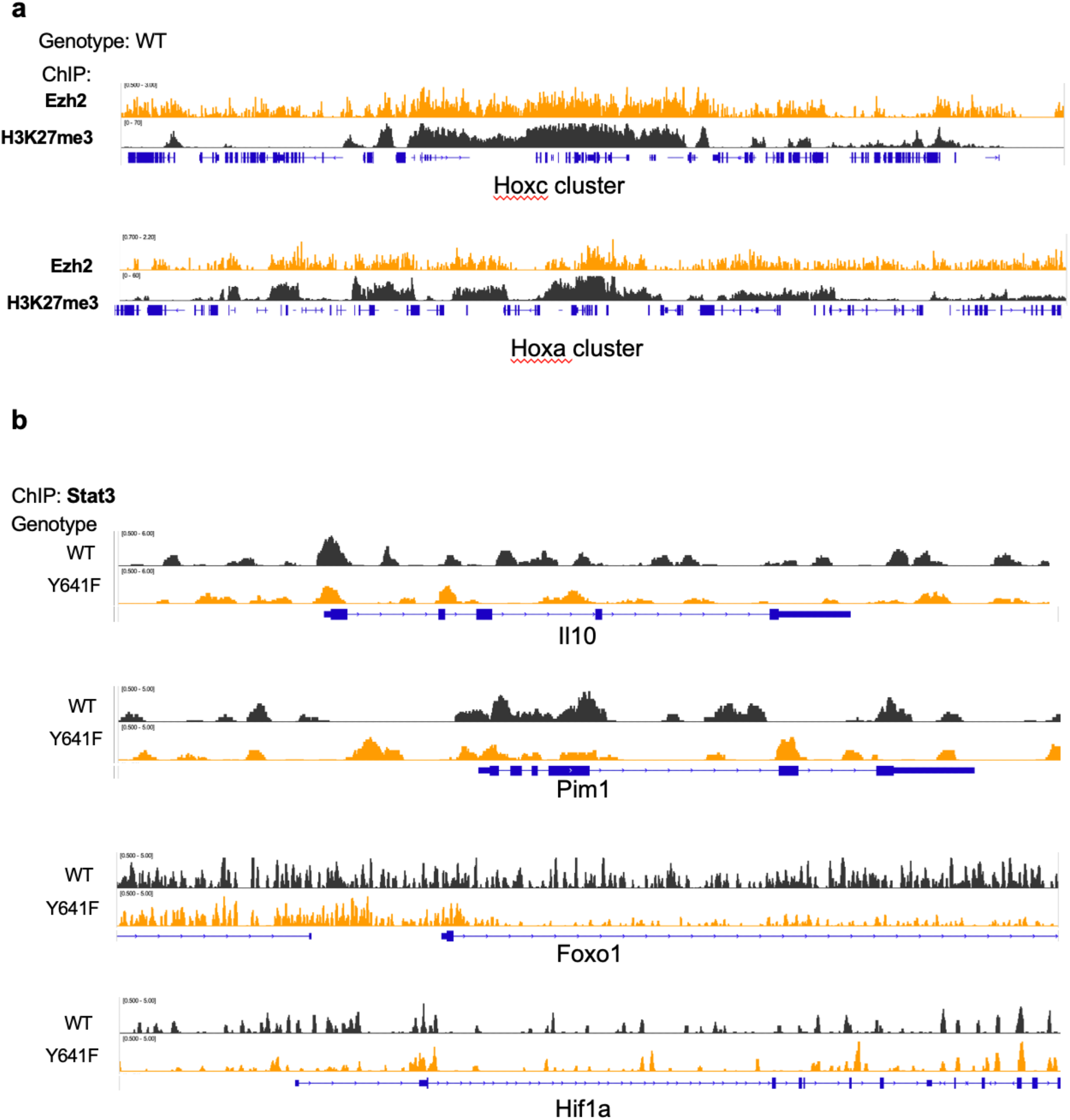
Representative loci of Ezh2 and Stat3 binding from ChIP-seq data. **(a)** Ezh2 ChIP-seq data showing binding at Hoxa and Hoxc clusters **(b)** Stat3 ChIP-seq data showing Stat3 peaks/binding at known Stat3 target genes: Il10, Pim1, Foxo1, Hif1a.

**Supp. Fig 4.**
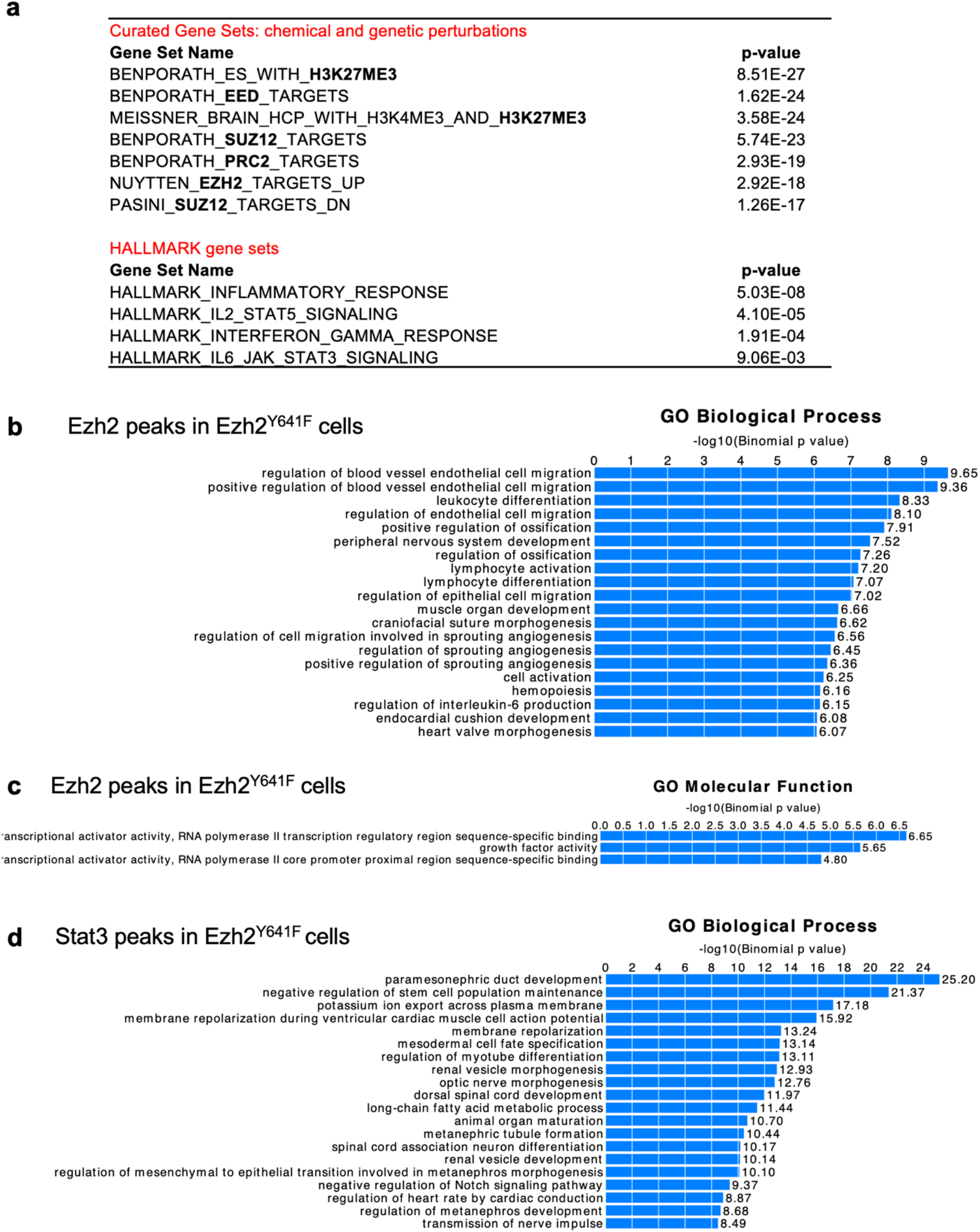
Functional annotation of differential ChIP-seq peaks in Ezh2^WT^ vs Ezh2^Y641F^ for Ezh2 and Stat3 proteins. **(a)** GSEA analysis of Ezh2 ChIP-seq peaks show enrichment for known PRC2 and EZH2 targets in *Ezh2*^WT^ and *Ezh2*^Y641F^ melanoma cells. **(b)** Gene Ontology Biological Process annotation for genes in proximity to Ezh2 ChIP-seq peaks unique to Ezh2^Y641F^ cells **(c)** Gene Ontology Molecular Function annotation for genes in proximity to Ezh2 ChIP-seq peaks unique to Ezh2^Y641F^ cells **(d)** Gene Ontology Biological Process annotation for genes in proximity to Stat3 ChIP-seq peaks unique to Ezh2^Y641F^ cells

